# Detection of neutralizing antibodies against arboviruses from liver homogenates

**DOI:** 10.1101/2024.05.28.596289

**Authors:** Thais Alkifeles Costa, Matheus Soares Arruda, Gabriela Fernanda Garcia Oliveira, Erik Vinicius de Sousa Reis, Anna Catarina Dias Soares Guimarães, Gabriel Dias Moreira, Nidia Esther Colquehuanca Arias, Marina do Vale Beirão, Nikos Vasilakis, Kathryn A. Hanley, Betânia Paiva Drumond

**Affiliations:** Laboratório de Vírus, Instituto de Ciências Biológicas, Universidade Federal de Minas Gerais, Belo Horizonte, MG, Brazil; Department of Pathology, University of Texas Medical Branch, Galveston, TX 77555-0609, USA; Center for Vector-Borne and Zoonotic Diseases, The University of Texas Medical Branch, Galveston, TX 77555-0609, USA; Institute for Human Infection and Immunity, University of Texas Medical Branch, Galveston, TX 77555-0610, USA; Department of Biology, New Mexico State University, Las Cruces, NM 88003-8801, USA

## Abstract

Yellow fever virus (YFV) circulates in a sylvatic cycle between non-human primates (NHPs) and arboreal mosquitoes in Brazil. Passive monitoring of ill or deceased NHPs is a key component of the Brazilian yellow fever (YF) surveillance program. Samples from NHPs carcasses are usually suitable for molecular tests but not for serological assays. As an alternative to the conventional plaque reduction neutralization test (PRNT) based on sera, we tested the utility of liver homogenates from experimentally infected (YFV, Mayaro virus [MAYV], chikungunya virus [CHIKV], or mock) mice to quantify PRNTs. Although homogenates from mock-infected mice showed a low level of nonspecific virus neutralization against YFV, MAYV or CHIKV, homogenates from YFV-, MAYV- and CHIKV-infected mice demonstrated significantly higher levels of virus neutralization compared to controls. Receiver operating characteristic (ROC) curves analyses were performed using the median neutralization values of three technical replicates for each infected group separately or collectively. Results showed scores ≥0.97 (95% CI ≥ 0.89-1.0) for the area under the curve at dilutions 1:20 to 1:80, suggesting that median virus neutralization values effectively differentiated YFV-, MAYV-, or CHIKV-infected groups from controls. Liver homogenates obtained from 25 NHPs carcasses (collected during the 2017 YF outbreak in Brazil) were also tested using the adapted PRNT as well as rapid anti-YFV IgM immunochromatographic tests. Neutralization activity was detected in six NHPs samples that were also positive by PCR and anti-YFV IgM tests and one sample that tested negative by PCR and IgM test. Our results demonstrate the feasibility of using liver homogenates as an alternative approach for serological investigation in viral epidemiologic surveillance.

## INTRODUCTION

Yellow fever virus (YFV) (*Flaviviridae, Orthoflavivirus*) is the causative agent of yellow fever (YF), a disease endemic in various tropical regions across Africa as well as Central and South America. Despite the existence of a highly effective vaccine, YF remains a significant global public health concern, as it is estimated that 80,000-200,000 YF cases occur annually in endemic regions, with a fatality rate ranging between 20% and 60% (WHO, 2024). Central and South America ended urban transmission of YFV in the 1950s though massive immunization campaigns, however the sylvatic cycle persists and serves as the source of periodic spillover to humans and epidemics throughout the region (reviewed by Sacchetto et al., 2020a; Silva et al., 2020).

In 2014, YFV reemerged in Midwest Brazil causing human cases and epizootics in the interface of the Brazilian Amazon, and Cerrado (Brazilian savannah) (Delatorre et al., 2019; Rezende et al., 2018; Ministério da Saúde, 2015; Ministério da Saúde, 2021). Subsequently, the virus spread to Southeastern Brazil, sparking a massive outbreak in Minas Gerais by December 2016, affecting thousands of humans and NHPs. Since then, YFV has spread into densely populated areas in Southeastern Brazil (Delatorre et al., 2019; Giovanetti et al., 2019; Figueiredo et al., 2020; Rezende et al., 2018; Ministério da Saúde, 2021; Silva et al., 2020), indicating that this region has suitable ecological and climatic conditions for YFV maintenance during the epidemic and interepidemic seasons (Rezende et al., 2018; Sacchetto et al., 2020b; Abreu et al., 2019; Possas et al., 2018). Between July 2014 and June 2023, 2,047 NHPs deaths; 2,289 human cases; and 780 human deaths (with a case fatality rate of 34%) caused by YFV were confirmed in the country (Giovanetti et al., 2023; PAHO, 2022; PAHO, 2023).

Neotropical NHPs are very susceptible to YFV infection and are considered key sentinels for outbreaks in the Americas (De Azevedo Fernandes et al., 2021; Sacchetto et al., 2020a; Sacchetto et al., 2020b; Silva et al., 2020). As reviewed by Silva and colleagues (2020) YFV infection in NHPs is similar to humans, causing a viscerotropic disease, with viral multiplication in various organs (liver, kidneys, bone marrow, spleen, and lymph nodes). Viremia is typically short-lived, ranging from three to seven days post-infection, but virus replication is still observed in the liver after that (Silva et al., 2020). One of the pillars of the Brazilian YF surveillance program is the investigation of NHPs epizootics. The early detection of YFV in NHPs can trigger control measures, such as human vaccination, thereby preventing outbreaks (Ministério da Saúde, 2017; Moreno et al., 2013; Silva et al., 2020). Epizootic surveillance primarily relies on passive monitoring of ill or deceased NHP of any species across Brazil. NHP carcasses constitute a primary source of biological samples for epizootic investigations, and laboratory tests are crucial for an accurate YF diagnosis (Ministério da Saúde, 2020).

The choice of laboratory tests to diagnose YF depends on the infection stage, the accessibility, and the quality of biological samples (PAHO, 2023). For laboratory diagnostic purposes, viral isolation and molecular tests are recommended during the infection phase. The sensitivity of molecular tests hinges on sample quality and the amount of viral RNA (PAHO, 2023). Various factors, such as temperature, organ degradation, and duration of postmortem exposure to weather conditions prior to sample collection, may influence the preservation status of NHP carcasses, potentially compromising the quality and integrity of viral RNA and hindering detection (Narat et al., 2018; Sacchetto et al., 2020a; Silva et al., 2023; Waggoner et al., 2018). After seroconversion, serological tests are preferred. IgM anti-YFV can be found in human serum samples six days post-exposure, and it may be detected up to four years in vaccinees (Reinhardt 1998; Gibney, 2012; Lindsey et al, 2018). As observed during the investigation of suspected YF human cases, the combined use of molecular and serological methods can enhance the sensitivity and specificity of the diagnosis (PAHO, 2023). However, the use of serological tests is frequently impractical due to difficulties or the impossibility of harvesting whole blood from carcasses (Kading et al., 2013; Waggoner et al., 2018). On the other hand, samples from solid tissues are easily harvested from carcasses and previous studies have demonstrated the potential for detecting antibodies using pig muscle and using blood samples of human cadavers (Gamble and Patrascu, 1996; Lai et al., 2022). In this context, utilizing solid organs obtained from NHPs carcasses could serve as a valuable strategy for investigating the presence of antibodies against YFV.

An alternative strategy for serological investigation of viral infections in samples from NHP carcasses, coupled with molecular tests, could significantly contribute to our understanding of the ecology and dynamics of various zoonotic viruses. Since many NHPs establish specific territories (reviewed by Silva et al., 2020), the detection of viral RNA or antibodies in local NHPs against specific viruses are good indicators of viral presence in a region. Besides YFV, NHPs act as significant arboviral reservoirs, creating the potential for spillover or spillback events for other orthoflaviviruses, as Zika virus (Terzian et al., 2018) and alphaviruses like Mayaro (MAYV) and chikungunya (CHIKV) in Brazil (Caicedo et al., 2021; Diagne et al., 2020; Moreira-Soto et al., 2018; Valentine et al., 2019). In our study, we investigated the presence of neutralizing antibodies against YFV, MAYV, and CHIKV using liver homogenates from experimentally infected mice as a substitute for serum. Subsequently, we applied the adapted protocol to investigate antibodies against YFV in liver samples from NHP carcasses collected during the YF epizootics in 2017 in Minas Gerais, Brazil.

## MATERIAL AND METHODS

### Cell line and viruses

Vero cells (ATCC CCL-81) were cultivated in Eagle’s minimum essential medium (MEM) (Cultilab, Brazil) supplemented with 5% fetal bovine serum (FBS) (Cultilab, Brazil) and antibiotics (200 U/mL penicillin [Cellofarm, Brazil]; 40 µg/mL streptomycin, [Sigma-Aldrich, Germany]; and 2 µg/mL amphotericin B [Cultilab, Brazil]) and incubated in a humidified 5% CO_2_ atmosphere at 37°C. Vero cells were used in cytotoxicity, PRNT, and virus titration assays.

The vaccine strain YFV-17DD Fiocruz (kindly provided by Dr. Pedro Augusto Alves from FIOCRUZ, Brazil), MAYV (BeAr20290, genotype L), and CHIKV (BHI1762H804917, genotype ECSA) (both kindly provided by Dr. Mauricio Lacerda Nogueira from FAMERP, Brazil) were used for mouse infection.

### Mouse strains, infection, and non-human primate samples

All the animal experiments received the approval of the Ethical Committee for Animal Experimentation of Universidade Federal de Minas Gerais (UFMG) (process no. 98/2017, approved in June - 29/2017; 33/2021, approved in March - 8/2021; 176/2021, approved in August - 23/2021).

Animal procedures were performed in mixed groups (males and females) of mice. For YFV experiments, IFNAR−/− mice (C57BL/6 background) aged 8 to 12 weeks were used. For MAYV and CHIKV experiments, C57BL/6 wild-type mice aged 3-4 weeks were used. Animals were housed in an animal care facility in individually ventilated cages at 23°C ± 2°C on a 12h/12h light/dark cycle with *ad libitum* access to water and food.

For each virus, mock (n = 6) and infected (n = 6) groups were utilized. One group infected with each virus was inoculated through the footpad route with 10 µL containing 1 × 10^3^ plaque-forming units (PFU) of YFV-17DD (Erickson and Pfeiffer, 2013); 1 × 10^5^ PFU of CHIKV (Hallengärd et al., 2014); or 1 × 10^5^ of MAYV (Mota et al., 2020). The mock-infected group was inoculated with the clarified supernatant of uninfected Vero cells (Fumagalli et al., 2021). Over 21 days, animals were observed for signs of infection. On day 21, mice were anesthetized with a solution of 100 mg/Kg of ketamine + 8-16 mg/Kg of xylazine per animal and were euthanized by cervical dislocation. Livers were harvested and stored at −80°C.

Liver samples collected from carcasses of free-living NHPs, during yellow fever outbreaks in 2017, in Minas Gerais, Brazil, were further utilized to obtain liver homogenates for serological testing. These samples were previously examined for YFV RNA by RT-qPCR (Sacchetto et al., 2020a).

### Sample processing

After euthanasia, liver samples were harvested from all mice (mock and infected groups), and separately processed to obtain a clarified liquid phase, referred to as a homogenate. Initially, liver samples were macerated utilizing four sterile beads in microtubes (Kasvi, USA) for 2 minutes in a bead beater with speed of 3,450 oscillations/min (Mini-Beadbeater-16, BioSpec Products, EUA), at a ratio of 50 mg of tissue to 200 µL of MEM without FBS. Subsequently, the samples were centrifuged at 16, 000 x g for 10 minutes, and the supernatant was transferred to new tubes (this clarification step was repeated five times). The homogenates were then incubated at 56°C without agitation (Thermomixer comfort, Eppendorf, Germany) for 30 minutes to heat inactivate the complement system. During this step, a clotting-like reaction altered the appearance of the homogenate samples. In response, two additional rounds of sample clarification (16,000 x g for 10 minutes) were performed. The final supernatants, referred to as liver homogenates, were stored at −20°C and utilized in cytotoxicity assays and for the viral neutralization tests.

Total RNA was extracted from approximately 30 mg of liver tissue from both infected and mock mice using the RNeasy Mini Kit (Qiagen, USA). Total RNA was then used to detect active infection by YFV (Domingo et al., 2012), MAYV (Naveca et al., 2017), or CHIKV (Edwards et al., 2007) via RT-qPCR, as previously described (Sacchetto et al., 2020a. Silva et al., 2023).

Liver samples obtained from NHPs were also utilized to produce liver homogenates following the previously described method. The resulting liver homogenate was diluted 1:10 in MEM without FBS, filtered with a 0.22 μm cellulose acetate membrane syringe microfilter (Filtrilo, Brazil), and then used in PRNT. Additionally, undiluted liver homogenate from NHP was employed in immunochromatographic tests for IgM anti-YFV (Febre amarela IgM EcoTeste, Ecodiagnóstica, Brazil).

### Cytotoxicity

To check whether liver homogenate could have a cytotoxic effect on Vero cells CCL-81, we performed 3-(4,5-dimethylthiazol-2-yl)-2,5-diphenyl-2H-tetrazolium bromide (MTT) reduction assay (Albarnaz et al., 2014). A total of 2.0 × 10^4^ Vero cells were seeded in each well of a 96-well plate and incubated for 24 h, at 37°C, and 5% CO_2_ atmosphere. Seventy microliters of liver homogenates were serially diluted (1:20 to 1:640) and added to Vero cells in 96-well plates in 200 µL of MEM 1% FBS, and incubated for two and five days, at 37°C, and 5% CO_2_ atmosphere. Afterward, the media was removed, and 28 µL of MTT solution (ThermoFisher Scientific, USA) in MEM (2 mg/mL) was added to each well. Cells were incubated at 37°C, and 5% CO_2_ atmosphere for 90 minutes protected from the light. Then, 130 µL of dimethyl sulfoxide solution (DMSO) was added to each well to solubilize formazan crystals. The plate, covered with aluminum foil, was agitated for 15 minutes. Finally, the absorbance at 540 nm was measured using a Multiskan GO Microplate spectrophotometer (Thermo Fisher Scientific, Waltham, Massachusetts, EUA), and the percentage decrease in cell viability was calculated. Simultaneously, under the same conditions, a crystal violet assay was performed to assess cell viability. In an identical plate, wells were washed with phosphate-buffered saline (PBS) three times, and 50 µL of 1% (w/v) crystal violet was added to each well and left to rest for 15 minutes. Plates were visually inspected regarding cell monolayer stability in the well. All conditions were tested in 12 well replicates.

### Adapted plaque reduction neutralization test (PRNT) with liver homogenate of experimentally infected mice

Liver homogenates from infected or mock mice, previously heat-inactivated and clarified, underwent triplicate testing by PRNT, in three technical replicates. Briefly, Vero CCL-81 cells (1.3 × 10^4^ cells per well) were seeded into 12-well plates and incubated at 37°C with a 5% CO_2_ atmosphere for 24 hours. Each virus was diluted using MEM 1% FBS, to achieve approximately 100-150 PFU per well. Liver homogenate samples (80 µL) were two-fold serially diluted (1:20 to 1:160) in MEM 1% FBS 1% HEPES buffer (4-[2-hydroxyethyl]-1-piperazineethanesulfonic acid).

Equal volumes of virus and diluted liver homogenate (1:10 to 1:160) were mixed, resulting in final dilutions of 1:20 to 1:320 of liver homogenates, and incubated at 37°C for 1 hour. Subsequently, 180 µL of the virus/liver homogenate mixture (1:20 to 1:160) was inoculated per well (in triplicate) onto Vero CCL-81 cells. Plates were incubated at 37°C for 1 hour at 5% CO_2_ atmosphere, with gentle rocking every 15 minutes. After incubation, the inoculum was removed, and each well was covered with 1 mL of 199 medium supplemented with 2% FBS and carboxymethyl cellulose (1% for YFV assay and 1.5% for CHIKV and MAYV assays) and plates were incubated at 37°C, 5% CO_2_ atmosphere. For YFV assay, plates were incubated for five days, while for CHIKV and MAYV assays, plates were incubated for two days, until the emergence of viral lysis plaques. Following incubation, plates were fixed with 3.7% formaldehyde solution for 1 hour and stained with 1% crystal violet solution for 30 minutes.

The serum used as positive controls were obtained from a biobank of samples from previously experimentally infected animals, as described above. Unfortunately, there were insufficient volumes of sera from the experimentally YFV-, MAYV-, CHIKV- or mock infected groups to be tested in parallel to liver homogenates. To assess whether liver homogenate could interfere with the neutralization activity, a second positive control, consisting of a 1:2 mixture of positive sera and liver homogenate from uninfected mice, was used. Liver homogenate from uninfected mice served as negative control. All samples and controls were tested in triplicate. Each plate included a well designated for cell control (no virus inoculation) and two wells designated for virus control.

Assay validity criteria included (i) the absence of PFU in the cell control well; (ii) neutralization by positive control sera, and no neutralization by negative control sera; (iii) the observed number of PFU for each triplicate (per each dilution of each mock or infected sample tested) in an experiment were within a 3-fold difference of the median PFU count (Dia et al., 2023; Timiryasova et al., 2013) and; (iv) the observation of 70 to 180 PFU in virus control wells. The median percentage of viral neutralization was estimated for each experiment comparing the median number of PFUs in virus wells to those in wells where the virus was incubated with liver homogenates, positive and negative controls. Three independent assays were conducted, with all PRNT assays completed by five researchers following the same standard assay procedures.

### Serological tests using liver homogenates of non-human primates

Filtered liver homogenates from NHPs were subjected to PRNT testing (dilution 1:20 to 640), following the previously described method. Undiluted liver homogenates were used to investigate the presence of IgM anti-YFV, using Febre amarela IgM EcoTest (EcoDiagnóstica, Brazil) following the manufacturer’s instructions. Briefly, 10 µL of liver homogenate were deposited in the specimen well of the cassette. Subsequently, 3 drops of the buffer provided by the manufacturer were added, and a 15-minute timer was set for result revelation.

### Data analyses

Statistical analyses were performed using GraphPrism version 8.0.2 for Windows (GraphPad Software, USA) or RStudio 4.3.2 (R Core Team, 2023, Wickham, 2016) with lme4 (Bates et al., 2015). For cytotoxicity analysis, linear regression was applied, with results with r^2^ > 0.9. Comparisons were made between the mock and infected groups for each homogenate dilution, and the differences were considered statistically significant when p ≤ 0.05. Generalized linear mixed models (GLMM) were employed to assess how neutralization responded to the status of each animal, designated by the infected or mock groups, by dilution (1:20 to 1:160), and by the interaction group/dilution. Individuals (animals) were treated as the random variable, while the explanatory variables included group (infected or mock) and dilution, and the interaction group/dilution. The response variable was virus neutralization. In different GLMM analyses, we used median neutralization values for both infected and mock groups (values per animal, per dilution, obtained from three independent assays (Table 1). All models were submitted to residual analysis to evaluate adequacy of error distribution (Crawley, 2013). Minimum adequate models were generated by stepwise omission of non-significant terms. Paired analyses (virus neutralization per group [mock or infected mice] per dilution) were conducted using GLMM with the group as the response variable and the animal as the random variable.

**Table 1:**
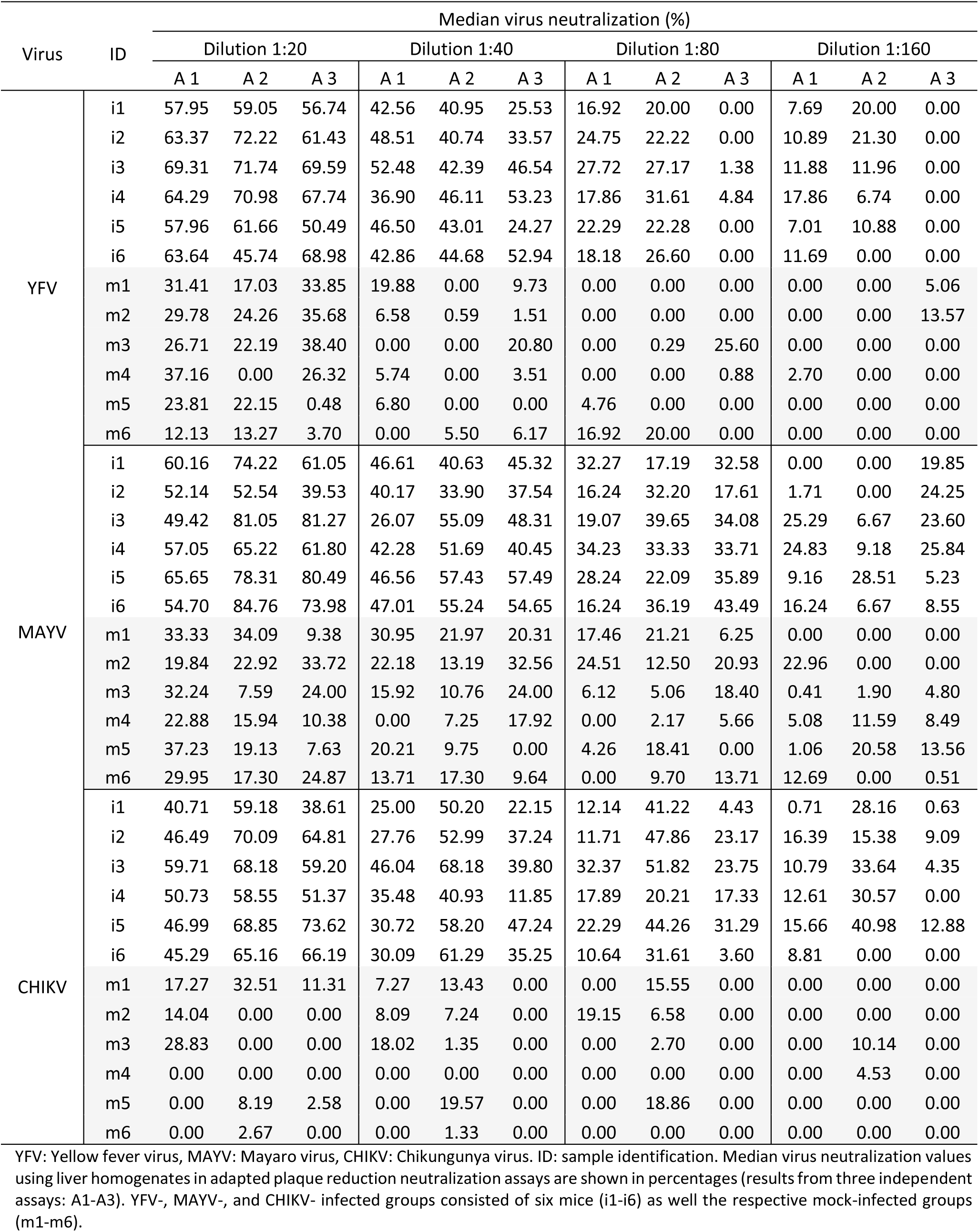
Median virus neutralization values (%) of YFV, MAYV, CHIKV using liver homogenates from experimentally infected mice in adapted plaque reduction neutralization tests.

| Virus | ID | Median virus neutralization (%) |  |  |  |  |  |  |  |  |  |  |  |
| --- | --- | --- | --- | --- | --- | --- | --- | --- | --- | --- | --- | --- | --- |
|  |  | Dilution 1:20 |  |  | Dilution 1:40 |  |  | Dilution 1:80 |  |  | Dilution 1:160 |  |  |
|  |  | A 1 | A 2 | A 3 | A 1 | A 2 | A 3 | A 1 | A 2 | A 3 | A 1 | A 2 | A 3 |
| YFV | i1 | 57.95 | 59.05 | 56.74 | 42.56 | 40.95 | 25.53 | 16.92 | 20.00 | 0.00 | 7.69 | 20.00 | 0.00 |
|  | i2 | 63.37 | 72.22 | 61.43 | 48.51 | 40.74 | 33.57 | 24.75 | 22.22 | 0.00 | 10.89 | 21.30 | 0.00 |
|  | i3 | 69.31 | 71.74 | 69.59 | 52.48 | 42.39 | 46.54 | 27.72 | 27.17 | 1.38 | 11.88 | 11.96 | 0.00 |
|  | i4 | 64.29 | 70.98 | 67.74 | 36.90 | 46.11 | 53.23 | 17.86 | 31.61 | 4.84 | 17.86 | 6.74 | 0.00 |
|  | i5 | 57.96 | 61.66 | 50.49 | 46.50 | 43.01 | 24.27 | 22.29 | 22.28 | 0.00 | 7.01 | 10.88 | 0.00 |
|  | i6 | 63.64 | 45.74 | 68.98 | 42.86 | 44.68 | 52.94 | 18.18 | 26.60 | 0.00 | 11.69 | 0.00 | 0.00 |
|  | m1 | 31.41 | 17.03 | 33.85 | 19.88 | 0.00 | 9.73 | 0.00 | 0.00 | 0.00 | 0.00 | 0.00 | 5.06 |
|  | m2 | 29.78 | 24.26 | 35.68 | 6.58 | 0.59 | 1.51 | 0.00 | 0.00 | 0.00 | 0.00 | 0.00 | 13.57 |
|  | m3 | 26.71 | 22.19 | 38.40 | 0.00 | 0.00 | 20.80 | 0.00 | 0.29 | 25.60 | 0.00 | 0.00 | 0.00 |
|  | m4 | 37.16 | 0.00 | 26.32 | 5.74 | 0.00 | 3.51 | 0.00 | 0.00 | 0.88 | 2.70 | 0.00 | 0.00 |
|  | m5 | 23.81 | 22.15 | 0.48 | 6.80 | 0.00 | 0.00 | 4.76 | 0.00 | 0.00 | 0.00 | 0.00 | 0.00 |
|  | m6 | 12.13 | 13.27 | 3.70 | 0.00 | 5.50 | 6.17 | 16.92 | 20.00 | 0.00 | 0.00 | 0.00 | 0.00 |
| MAYV | i1 | 60.16 | 74.22 | 61.05 | 46.61 | 40.63 | 45.32 | 32.27 | 17.19 | 32.58 | 0.00 | 0.00 | 19.85 |
|  | i2 | 52.14 | 52.54 | 39.53 | 40.17 | 33.90 | 37.54 | 16.24 | 32.20 | 17.61 | 1.71 | 0.00 | 24.25 |
|  | i3 | 49.42 | 81.05 | 81.27 | 26.07 | 55.09 | 48.31 | 19.07 | 39.65 | 34.08 | 25.29 | 6.67 | 23.60 |
|  | i4 | 57.05 | 65.22 | 61.80 | 42.28 | 51.69 | 40.45 | 34.23 | 33.33 | 33.71 | 24.83 | 9.18 | 25.84 |
|  | i5 | 65.65 | 78.31 | 80.49 | 46.56 | 57.43 | 57.49 | 28.24 | 22.09 | 35.89 | 9.16 | 28.51 | 5.23 |
|  | i6 | 54.70 | 84.76 | 73.98 | 47.01 | 55.24 | 54.65 | 16.24 | 36.19 | 43.49 | 16.24 | 6.67 | 8.55 |
|  | m1 | 33.33 | 34.09 | 9.38 | 30.95 | 21.97 | 20.31 | 17.46 | 21.21 | 6.25 | 0.00 | 0.00 | 0.00 |
|  | m2 | 19.84 | 22.92 | 33.72 | 22.18 | 13.19 | 32.56 | 24.51 | 12.50 | 20.93 | 22.96 | 0.00 | 0.00 |
|  | m3 | 32.24 | 7.59 | 24.00 | 15.92 | 10.76 | 24.00 | 6.12 | 5.06 | 18.40 | 0.41 | 1.90 | 4.80 |
|  | m4 | 22.88 | 15.94 | 10.38 | 0.00 | 7.25 | 17.92 | 0.00 | 2.17 | 5.66 | 5.08 | 11.59 | 8.49 |
|  | m5 | 37.23 | 19.13 | 7.63 | 20.21 | 9.75 | 0.00 | 4.26 | 18.41 | 0.00 | 1.06 | 20.58 | 13.56 |
|  | m6 | 29.95 | 17.30 | 24.87 | 13.71 | 17.30 | 9.64 | 0.00 | 9.70 | 13.71 | 12.69 | 0.00 | 0.51 |
| CHIKV | i1 | 40.71 | 59.18 | 38.61 | 25.00 | 50.20 | 22.15 | 12.14 | 41.22 | 4.43 | 0.71 | 28.16 | 0.63 |
|  | i2 | 46.49 | 70.09 | 64.81 | 27.76 | 52.99 | 37.24 | 11.71 | 47.86 | 23.17 | 16.39 | 15.38 | 9.09 |
|  | i3 | 59.71 | 68.18 | 59.20 | 46.04 | 68.18 | 39.80 | 32.37 | 51.82 | 23.75 | 10.79 | 33.64 | 4.35 |
|  | i4 | 50.73 | 58.55 | 51.37 | 35.48 | 40.93 | 11.85 | 17.89 | 20.21 | 17.33 | 12.61 | 30.57 | 0.00 |
|  | i5 | 46.99 | 68.85 | 73.62 | 30.72 | 58.20 | 47.24 | 22.29 | 44.26 | 31.29 | 15.66 | 40.98 | 12.88 |
|  | i6 | 45.29 | 65.16 | 66.19 | 30.09 | 61.29 | 35.25 | 10.64 | 31.61 | 3.60 | 8.81 | 0.00 | 0.00 |
|  | m1 | 17.27 | 32.51 | 11.31 | 7.27 | 13.43 | 0.00 | 0.00 | 15.55 | 0.00 | 0.00 | 0.00 | 0.00 |
|  | m2 | 14.04 | 0.00 | 0.00 | 8.09 | 7.24 | 0.00 | 19.15 | 6.58 | 0.00 | 0.00 | 0.00 | 0.00 |
|  | m3 | 28.83 | 0.00 | 0.00 | 18.02 | 1.35 | 0.00 | 0.00 | 2.70 | 0.00 | 0.00 | 10.14 | 0.00 |
|  | m4 | 0.00 | 0.00 | 0.00 | 0.00 | 0.00 | 0.00 | 0.00 | 0.00 | 0.00 | 0.00 | 4.53 | 0.00 |
|  | m5 | 0.00 | 8.19 | 2.58 | 0.00 | 19.57 | 0.00 | 0.00 | 18.86 | 0.00 | 0.00 | 0.00 | 0.00 |
|  | m6 | 0.00 | 2.67 | 0.00 | 0.00 | 1.33 | 0.00 | 0.00 | 0.00 | 0.00 | 0.00 | 0.00 | 0.00 |

YFV: Yellow fever virus, MAYV: Mayaro virus, CHIKV: Chikungunya virus. ID: sample identification. Median virus neutralization values using liver homogenates in adapted plaque reduction neutralization assays are shown in percentages (results from three independent assays: A1-A3). YFV-, MAYV-, and CHIKV- infected groups consisted of six mice (i1-i6) as well the respective mock-infected groups (m1-m6).

GLMM were also used to assess how neutralization of mock or infected animals responded to the virus used in each assay (YFV, MAYV and CHIKV), designated by virus, by dilution (1:20 to 1:160), and by the interaction virus/dilution. Analyses were carried out considering mock and infected groups separately. Individuals were treated as the random variable, while the explanatory variables included virus (YFV, MAYV or CHIKV), dilution, and the interaction virus/dilution. The response variable was median virus neutralization (values per animal, per dilution), obtained from three independent assays (Table 1). Post-hoc analyses for pairwise comparisons were performed, with Tukey correction, using the glht function in Multcomp package (Hothorn et al, 2008) and the differences were considered statistically significant when p ≤ 0.05.

Neutralization percentage data were subjected to Receiver operating characteristic (ROC) curve analyses performed using GraphPad Prism version 8.0.2 for Windows (GraphPad Software, USA) using the Wilson/Brown method (Wilson, 1927; Brown, et al., 2001). For these analyses, median values (Table 1) were compared by dilution for YFV, MAYV, and CHIKV separately. Additionally, we performed the same analysis using median neutralization values for the three viruses combined (Table 1), also grouped by dilution. Moreover, we compared the lowest median neutralization values obtained from infected animal samples with the highest neutralization values obtained from mock-infected samples for each dilution, to investigate the occurrence of overlapping neutralization amongst both groups (Table 1).

## RESULTS AND DISCUSSION

### Neutralizing activity of liver homogenates of experimentally infected mice against YFV, MAYV, and CHIKV

To test whether virus-neutralizing antibodies could be detected from the livers of animal carcasses, we employed liver homogenates obtained from mice experimentally infected with YFV, MAYV, CHIKV, or subjected to mock infection for performing PRNTs. The choice of the liver was motivated by its role as a significant blood reservoir, receiving 25% of the cardiac output while constituting only 2.5% of the body weight (Lautt et al., 2009). We infected mice with non-lethal doses of YFV, MAYV, and CHIKV, which we evaluated for infection signs up to 21 dpi, and at this date, we euthanized animals and harvested the liver without venipuncture. All (YFV-, MAYV-, and CHIKV-) infected animals presented general signs of infection, such as ruffled fur and hunched posture. For CHIKV- and MAYV-infected mice, paw edema until 3 dpi was also observed. Mock-infected animals presented no reaction to mock components. At 21 dpi, all liver samples were negative for the presence of RNA for YFV, MAYV, or CHIKV by RT-qPCR.

Cytotoxicity of liver homogenates for Vero cells was then assessed. Liver homogenates of C57BL/6 (n = 3) and IFNAR −/− (n = 4) animals were incubated in Vero CCL-81 cells for two and five days, respectively, and cytotoxicity was evaluated by an MTT assay. At two days of incubation (Fig. S1), at the 1:20 dilution, the mean cell viability was 80.1%, and above that (1:40 and beyond) the viability was > 80%. At five days of incubation (Fig. S1), at the 1:20 dilution, the mean cell viability was 75.97%, and above that (1:40 and beyond) the viability was > 80% (Fig. S1).

Next, the adapted PRNT was performed, with serum being replaced by the clarified liver homogenate. Liver homogenates from both mock and YFV-infected animals were evaluated in neutralization assays against YFV. Median virus neutralization of 63.50% (45.74 – 72.22%) at dilution 1:20; 42.93% (24.27 – 53.23%) at dilution 1:40; 19.09% (0 – 31.61%) at dilution 1:80; and 7.35% (0 – 21.30%) at dilution 1:160 were observed for the YFV-infected group across three independent assays (Table 1). The mock group displayed lower median neutralization values of 24.03% (0.0 – 38.40%) at dilution 1:20; 2.51% (0.0 – 20.80%) at dilution 1:40; 0% (0.0 – 25.60%) at dilution 1:80; and 0.0% (0.0 – 13.57%) at dilution 1:160 (Table 1). The positive control sera from infected mice exhibited median neutralization values of 96.50% (85.63 – 99.11%) at dilution 1:20; 84.78% (82.04 – 92.86%) at dilution 1:40; 77.62% (63.52 – 79.46%) at dilution 1:80; and 50.0% (32.08 - 50.72%) at dilution 1:160. The positive control 2, consisting of a 1:2 mixture of liver homogenate from mock-infected mice sera from infected mice, exhibited median neutralization values of 95.69% (91.84 – 97.47%) at dilution 1:20; 85.11% (79.59 – 89.87%) at dilution 1:40; 56.90% (53.74 – 73.42%) at dilution 1:80; and 33.33% (14.89 – 50.86%) at dilution 1:160. These results suggest that the liver homogenate does not prevent YFV neutralization.

Liver homogenates of mock or MAYV-infected animals were tested against MAYV in adapted PRNT. For the MAYV-infected group, the median virus neutralization values ranged from 63.51% (39.53 – 84.76%) at dilution 1:20, to 46.59% (26.07 – 57.49%) at dilution 1:40; 32.43% (16.24 – 43.49%) at dilution 1:80; and 9.17% (0.0 – 28.51%) at dilution 1:160, considering three independent assays (Table 1). The mock group showed lower median neutralization values of 22.90% (7.59 – 37.23%) at dilution 1:20; 16.61% (0 – 32.56%) at dilution 1:40; 7.98% (0 – 24.51%) at dilution 1:80; and finally, 1.48% (0.0 – 22.96%) at dilution 1:160 (Table 1). The positive control sera from infected mice, exhibited median neutralization values of 85.37% (84.0 - 91.55%) at dilution 1:20; 79.93% (78.20 – 81.69%) at dilution 1:40; 66.49% (52.11 – 76.77%) at dilution 1:80; and 49.31% (45.78 – 58.71%) at dilution 1:160. The positive control 2, consisting of a 1:2 mixture of liver homogenate from mock-infected mice sera from infected mice, exhibited median neutralization values of 85.87% (78.53 – 88.13%) at dilution 1:20; 75.33% (73.12 – 79.0%) at dilution 1:40; 65.43% (54.44 – 74.43%) at dilution 1:80; and 42.33% (36.73 – 63.84%) at dilution 1:160. The results also suggest that the liver homogenate does not prevent MAYV neutralization.

Liver homogenates of mock- or CHIKV-infected animals were challenged against CHIKV in PRNT. For CHIKV-infected group, the median virus neutralization values ranged from 59.19% (38.61 – 73.62%) at dilution 1:20 to 38.52% (11.85 – 68.18%) at dilution 1:40; 22.73% (3.60 – 51.82%) at dilution 1:80; and 11.70% (0.0 – 40.98%) at dilution 1:160, considering three independent assays (Table 1). The mock group showed median neutralization values lower of 0.0% (0.0 – 32.51%) at dilution 1:20; 0.0% (0.0 – 19.57%) at dilution 1:40; 0.0% (0.0 – 19.15%) at dilution 1:80; and finally, 0% (0.0 – 10.14%) at dilution 1:160 (Table 1). The positive control sera from infected mice, exhibited median neutralization values of dilutions of 85.43% (58.02 – 89.89%) at dilution 1:20; 81.82% (58.02 – 94.95%) at dilution 1:40; 75.23% (60.75 – 94.22%) at dilution 1:80; and 69.23% (34.34 – 94.12%) at dilution 1:160. The positive control 2, consisting of a 1:2 mixture of liver homogenate from mock-infected mice sera from infected mice, exhibited median neutralization values of 89.83% (84.54 – 90.77%) at dilution 1:20; 84.90% (77.52 – 91.03%) at dilution 1:40; 79.91% (70.85 –85.90%) at dilution 1:80; and 61.35% (51.38 – 78.21%) at dilution 1:160. Similarly to YFV and MAYV, neutralization of CHIKV was not prevented by the liver homogenate.

Generalized linear mixed model analyses revealed significant differences in average median neutralization regarding the groups (mock or infected), dilution, and the interaction group/dilution (p < 0.001, Table 2, Fig.1). The results indicated that as the dilution increased the difference between infected and mock groups decreased. This pattern could be explained by the dilution of specific neutralizing antibodies in liver homogenates of infected mice, in contrast to their absence in the mock group (Table 2, Fig 1). Similarly, pairwise analyses of the interaction group/dilution demonstrated that the difference in average neutralization between the YFV-, MAYV-, and CHIKV-infected groups and their respective mock groups decreased up to dilution 1:80 (see the bold numbers in Table 3). Differences were observed between infected and their respective mock groups when comparing the average median viral neutralization values at dilutions 1:20 to 1:80 with each other and with dilution 1:160 (p< 0.01, Table 3). With few exceptions, the median neutralization values observed at dilution 1:160 in infected mice were not significantly different from the values observed for mock mice at dilutions 1:20 to 1:160 (except for YFV-infected at dilution 1:160 vs mock at dilution 1:20, and CHIKV-infected and all viruses at dilution 1:160 vs mock at dilution 1:160) (Table 3). The differences between infected and mock groups could be attributed to the presence of specific neutralizing antibodies in the liver homogenates of infected mice in contrast to their absence in mock mice.

**Figure 1:**
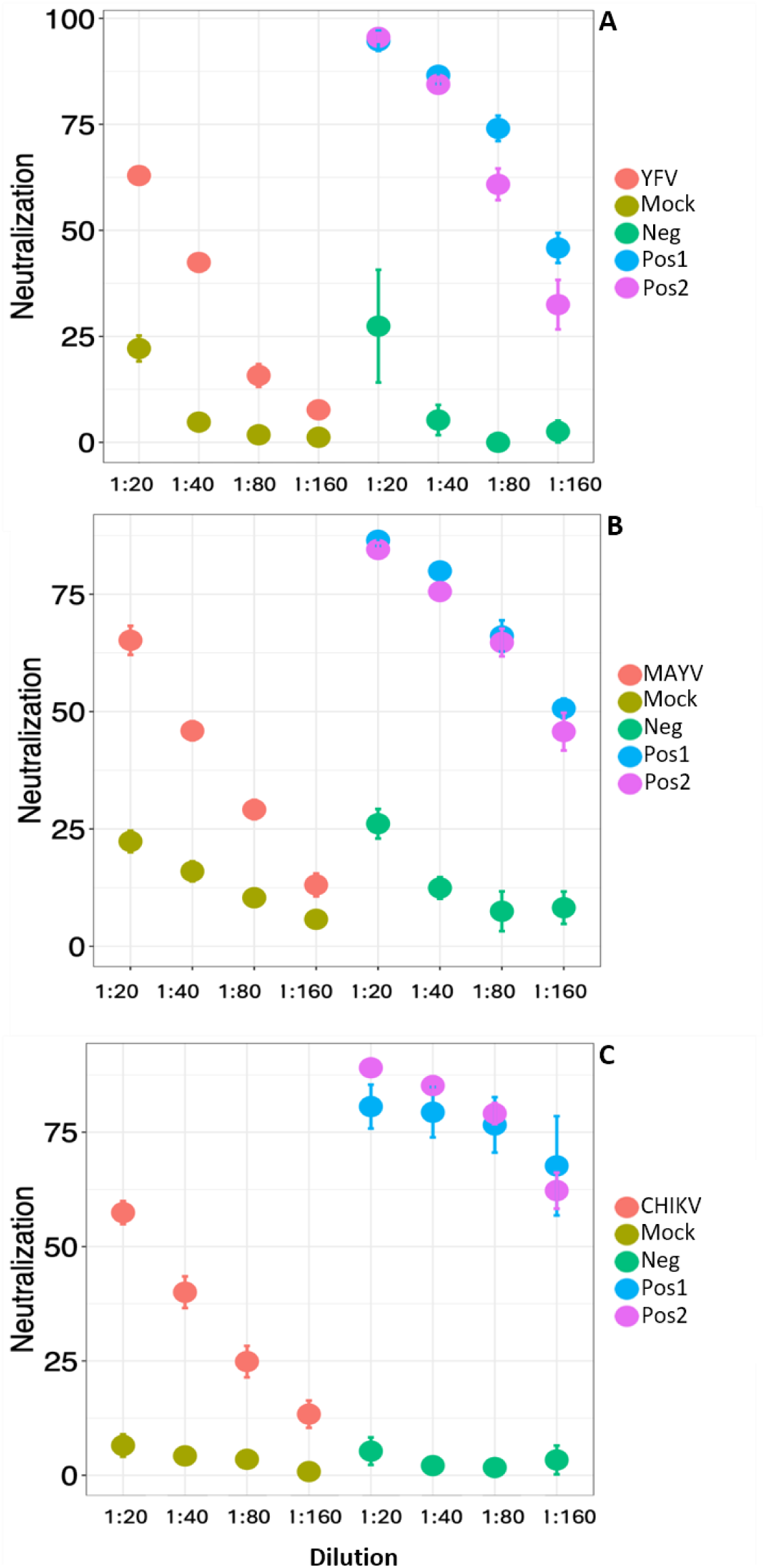
Average median virus neutralization values of YFV (A), MAYV (B), CHIKV (C) per dilution, using liver homogenates from experimentally infected mice in adapted plaque reduction neutralization tests. A, B and C respectively display average median neutralization values (in percentage) obtained from the experiments carried out with samples from YFV-, MAYV- and CHIKV-infected groups, and their respective mock-groups. Bars represent standard deviation. YFV-, MAYV-, and CHIKV-infected groups consisted of six mice each as well the respective mock-infected groups. The negative control is liver homogenate obtained from uninfected animals (neg), the positive control is sera from experimentally infected animal (pos 1), and the second positive control is composed of a mixture (1:2) of liver homogenate of uninfected animal and positive serum (pos2).

**Table 2:** Results of generalized linear mixed models analysis of the response of average median virus neutralization to groups infected and mock, dilution and group/dilution in adapted plaque reduction neutralization tests using liver homogenates of experimentally infected mice.

| Response variable | Explanatory variables | F | p |
| --- | --- | --- | --- |
| YFV neutralization (%) | group (YFV infected/mock) | 309.13 | < 0.001 |
|  | dilution | 154.13 | < 0.001 |
|  | group/dilution | 37.45 | < 0.001 |
| MAYV neutralization (%) | group (MAYV infected/mock) | 204.41 | < 0.001 |
|  | dilution | 72.97 | < 0.001 |
|  | group/dilution | 19.31 | < 0.001 |
| CHIKV neutralization (%) | group (CHIKV infected/mock) | 283.58 | < 0.001 |
|  | dilution | 35.66 | < 0.001 |
|  | group/dilution | 22.07 | < 0.001 |
| YFV + MAYV + CHIKV neutralization (%) | group (infected/mock) | 590.48 | < 0.001 |
|  | dilution | 168.85 | < 0.001 |
|  | group/dilution | 55.23 | < 0.001 |

YFV: Yellow fever virus, MAYV: Mayaro virus, CHIKV: Chikungunya virus. YFV-, MAYV-, and CHIKV-infected groups consisted of six mice each as well the respective mock-infected groups.

**Table 3.** – Pairwise comparison of average median virus neutralization per group/dilution in adapted plaque reduction neutralization tests using liver homogenates of experimentally infected mice.

| virus group/dilution |  | Mock (YFV) |  |  |  | YFV-infected |  |  |
| --- | --- | --- | --- | --- | --- | --- | --- | --- |
|  |  | 1:20 | 1:40 | 1:80 | 1:160 | 1:40 | 1:80 | 1:160 |
| YFV-Infected | 1:20 | **<0.01\*** | <0.01* | <0.01* | <0.01* | <0.01* | <0.01* | <0.01* |
|  | 1:40 | <0.01* | **<0.01\*** | <0.01* | <0.01* |  | <0.01* | <0.01* |
|  | 1:80 | <0.01* | <0.01* | **<0.01\*** | <0.01* |  |  | 0.067 |
|  | 1:160 | <0.01* | 0.970 | 0.397 | **0.272** |  |  |  |
| Mock (YFV) | 1:40 | <0.01* |  |  |  |  |  |  |
|  | 1:80 | <0.01* | 0.964 |  |  |  |  |  |
|  | 1:160 | <0.01* | 0.908 | 0.888 |  |  |  |  |

| virus group/dilution |  | Mock (MAYV) |  |  |  | MAYV-infected |  |  |
| --- | --- | --- | --- | --- | --- | --- | --- | --- |
|  |  | 1:20 | 1:40 | 1:80 | 1:160 | 1:40 | 1:80 | 1:160 |
| MAYV - Infected | 1:20 | **<0.01\*** | <0.01* | <0.01* | <0.01* | <0.01* | <0.01* | <0.01* |
|  | 1:40 | <0.01* | **<0.01\*** | <0.01* | <0.01* |  | <0.01* | <0.01* |
|  | 1:80 | 0.511 | <0.01* | **<0.01\*** | <0.01* |  |  | <0.01* |
|  | 1:160 | 0.128 | 0.991 | 0.994 | **0.402** |  |  |  |
| Mock (MAYV) | 1:40 | 0.589 |  |  |  |  |  |  |
|  | 1:80 | 0.012* | 0.734 |  |  |  |  |  |
|  | 1:160 | 0.01* | 0.062 | 0.888 |  |  |  |  |

| virus group/dilution |  | Mock (CHIKV) |  |  |  | CHIKV-infected |  |  |
| --- | --- | --- | --- | --- | --- | --- | --- | --- |
|  |  | 1:20 | 1:40 | 1:80 | 1:160 | 1:40 | 1:80 | 1:160 |
| CHIKV - Infected | 1:20 | **<0.001\*** | <0.001* | <0.001* | <0.001* | <0.001* | <0.001* | <0.001* |
|  | 1:40 | <0.001* | **<0.001\*** | <0.001* | <0.001* |  | <0.001* | <0.001* |
|  | 1:80 | <0.001* | <0.001* | **<0.001\*** | <0.001* |  |  | 0.029* |
|  | 1:160 | 0.543 | 0.175 | 0.106 | **0.011\*** |  |  |  |
| Mock (CHIKV) | 1:40 | 0.998 |  |  |  |  |  |  |
|  | 1:80 | 0.990 | 1.000 |  |  |  |  |  |
|  | 1:160 | 0.754 | 0.980 | 0.996 |  |  |  |  |

| virus group/dilution |  | Mock (all viruses) |  |  |  | all viruses-infected |  |  |
| --- | --- | --- | --- | --- | --- | --- | --- | --- |
|  |  | 1:20 | 1:40 | 1:80 | 1:160 | 1:40 | 1:80 | 1:160 |
| all viruses-infected | 1:20 | **<0.001\*** | <0.001* | <0.001* | <0.001* | <0.001* | <0.001* | <0.001* |
|  | 1:40 | <0.001* | **<0.001\*** | <0.001* | <0.001* |  | <0.001* | <0.001* |
|  | 1:80 | 0.070 | <0.001* | **<0.001\*** | <0.001* |  |  | <0.001* |
|  | 1:160 | 0.184 | 0.871 | 0.086 | **0.001\*** |  |  |  |
| mock-(all viruses) | 1:40 | 0.003* |  |  |  |  |  |  |
|  | 1:80 | <0.001* | 0.832 |  |  |  |  |  |
|  | 1:160 | <0.001* | 0.138 | 0.931 |  |  |  |  |

YFV: Yellow fever virus, MAYV: Mayaro virus, CHIKV: Chikungunya virus. Post-hoc analyses of generalized linear mixed models, with Tukey correction. The asterisks indicate statistical significance, and the bold numbers indicate the comparison between groups in the respective dilution.

The average median neutralization differed between serial dilutions (p<0.029) (e.g. 1:20 vs 1:40, 1:20 vs 1:80 and so on) within each infected group (YFV, MAYV, CHIKV, and all viruses combined) (Table 3) with a single exception within YFV-group (dilution 1:80 vs. 1:160, Table 3). The variations in viral neutralization observed among serial dilutions of the infected group may be linked to the presence of specific neutralizing antibodies in the liver homogenates of infected mice, which respond accordingly as the dilution increases. On the other hand, the average median neutralization values did not differ between serial dilutions (1:40 to 1:160) within each mock group (tested against YFV, MAYV, CHIKV, and for all viruses combined). Differences in the average median neutralization values for mock groups were only observed when the lowest dilution, 1:20, was compared to higher dilutions (1:40 to 1:160 for YFV and for all viruses combined, and 1:80 to 1:160 for MAYV) (Table 3). These results indicate that in the dilution 1:20, a higher nonspecific neutralization could be observed, but as the dilution factor increased no effect on viral neutralization was observed and the reduction on PFUs observed for the mock group could be related to nonspecific neutralization.

To assess whether the average neutralization values could differ depending on the virus used in each assay, we separated mock from infected groups and performed GLMM analyses considering virus, dilution, and the interaction virus/dilution. When mock groups were compared to each other, significant differences in average median neutralization regarding the virus, the dilution, and the interaction virus/dilution (p<0.001) were observed (Table 4). The results indicated that as the dilution increased the difference among types of viruses decreased. This could be explained by the dilution of components of liver homogenate leading to nonspecific neutralization when using mock samples. In addition, post-hoc pairwise comparisons demonstrated that the mock group tested against CHIKV exhibited lower values of nonspecific neutralization compared to the mock groups tested against YFV and MAYV at dilution 1:20 (p <0.01, Fig.S2). Conversely, at dilutions 1:40 to 1:160, mock animals tested against CHIKV and YFV presented similar neutralization values compared to each other (p≥0.97, Fig.S2). While YFV and MAYV had similar neutralization values at dilution 1:20, the effect of nonspecific neutralization was higher for MAYV compared to YFV at dilutions 1:40 and 1:80 (p≤0.03, Fig.S2), and compared to CHIKV at dilutions 1:20 and 1:40 (p≤0.01, Fig.S2). The results indicate that nonspecific neutralization could vary depending on the virus investigated and the dilution and that MAYV had a higher influence of non-specific neutralization in higher dilutions compared to YFV and CHIKV.

**Table 4:** Results of generalized linear mixed models’ analysis of the response of average median virus neutralization to virus, dilution and virus/dilution in adapted plaque reduction neutralization tests using liver homogenates of experimentally infected or mock mice.

| Group | Response variable | Explanatory variables | F | p |
| --- | --- | --- | --- | --- |
| Mock<sup>a</sup> | neutralization (%) | virus | 31.8031 | < 0.001* |
|  |  | dilution | 17.2953 | < 0.001* |
|  |  | virus/dilution | 5.7768 | < 0.001* |
| Infected<sup>b</sup> | neutralization (%) | virus | 6.1071 | <0.001* |
|  |  | dilution | 193.7954 | <0.001* |
|  |  | virus/dilution | 2.1187 | 0.053 |
| Infected<sup>c</sup> | neutralization (%) | virus | 5.8987 | 0.003* |
|  |  | dilution | 193.7955 | < 0.001* |

<sup>a</sup>complete significant model, <sup>b</sup>complete non-significant model, <sup>c</sup>minimal adequate model.

When infected groups were compared to each other, GLMM analyses did not reveal significant differences regarding to the interaction virus/dilution (p =0.053, Table 4), indicating that the neutralization observed for one specific virus at a certain dilution is independent from the neutralization observed for another virus at a given dilution, as expected for virus-specific neutralization. The type of virus (p=0.003) and dilution (p<0.001) both significantly affected the average median neutralization (Table 4). Pairwise analyses indicated differences between all serial dilutions, with the decrease of average median neutralization values towards higher dilutions (p< 1.0^-6^, Fig.S3 A and B). Regardless of the dilution, MAYV presented higher median neutralization values compared to YFV (p<0.01) and CHIKV (p=0.044), while no differences were observed between CHIKV and YFV (p=0.62, Fig.S3 C and D). The higher neutralization observed for MAYV-infected group, compared to YFV- and CHIKV-infected groups could be related to the higher level of nonspecific neutralization observed for MAYV (as demonstrated in the mock group) or to intrinsic properties of MAYV neutralization. The results showed that when using liver homogenates, we can observe virus neutralization for all viruses investigated. The effect of the type of virus on neutralization is consistent across dilutions, and the effect of dilution on neutralization is consistent across types of viruses.

The data revealed a baseline virus neutralization in both mock and infected groups, potentially attributable to the composition of the liver homogenate. Even after multiple rounds of clarification, it is plausible that some residual cellular or tissue debris persists as particulate organic matter in the liver homogenate. This particulate organic matter might interfere with the viral infection process by adsorbing to viral particles or the cell monolayer, leading to nonspecific virus neutralization. Despite the nonspecific virus neutralization observed in mock groups, infected mice exhibited higher virus neutralization compared to mock mice, which we attribute to the presence of neutralizing antibodies enacting specific virus neutralization.

ROC curve analyses were conducted utilizing the median neutralization values from three independent assays (Table 1) for each infected group separately (YFV, MAYV, or CHIKV) and collectively (YFV+MAYV+CHIKV), alongside their corresponding mock groups. AUC scores ≥0.97 (95%CI ≥0.89 – 1.0, p<0.0065) indicated that the median virus neutralization values effectively differentiated YFV-, MAYV- or CHIKV-infected groups from their respective mock groups at dilutions 1:20 to 1:80 (Table 5). Moreover, ROC curve analysis encompassing all infected groups versus all mock groups per dilution also demonstrated robust AUC scores ≥ 0.97 (95% CI ≥ 0.92 – 1.0, p< 0.0001), effectively discriminating both groups at dilutions 1:20 to 1:80 (Table 5). The ROC curve analyzes in agreement with GLMM results, confirmed that median virus neutralization values successfully discriminated between infected and mock groups across all dilutions.

**Table 5:** Receiver operating characteristic (ROC) curve analyses for median neutralization values of YFV-, MAYV, CHIKV-infected groups compared to their respective mock groups.

| Virus<br>(sample<br>size) | dil | Median neutralization<br>values infected x mock |  | Cutoff | Se (95% CI) | Sp (95% CI) |
| --- | --- | --- | --- | --- | --- | --- |
|  |  | AUC (95% CI) | p value |  |  |  |
| YFV<br>(n=12) | 1:20 | 1.0 (1.0-1.0) | 0.0039* | 57.96 | 0.83 (0.44-0.99) | 1.0 (0.61 – 1.0) |
|  | 1:40 | 1.0 (1.0-1.0) | 0.0039* | 40.85 | 0.83 (0.44-0.99) | 1.0 (0.61 – 1.0) |
|  | 1:80 | 1.0 (1.0-1.0) | 0.0039* | 17.39 | 0.83 (0.44-0.99) | 1.0 (0.61 – 1.0) |
|  | 1:160 | 0.92 (0.73-1.0) | 0.0163* |  |  |  |
| MAYV<br>(n=12) | 1:20 | 1.0 (1.0-1.0) | 0.0039* | 56.59 | 0.83 (0.44-0.99) | 1.0 (0.61 -1.0) |
|  | 1:40 | 1.0 (1.0-1.0) | 0.0039* | 39.91 | 0.83 (0.44-0.99) | 1.0 (0.61-1.0) |
|  | 1:80 | 0.97 (0.89-1.0) | 0.0065* | 24.59 | 0.83 (0.44-0.99) | 1.0 (0.61-1.0) |
|  | 1:160 | 0.72 (0.42-1) | 0.20 |  |  |  |
| CHIKV<br>(n=12) | 1:20 | 1.0 (1.0-1.0) | 0.0039* | 46.04 | 0.83 (0.44-0.99) | 1.0 (0.61-1.0) |
|  | 1:40 | 1.0 (1.0 –1.0) | 0.0039* | 30.13 | 0.83 (0.44-0.99) | 1.0 (0.61-1.0) |
|  | 1:80 | 1.0 (1.0 –1.0) | 0.0039* | 11.39 | 0.83 (0.44-0.99) | 1.0 (0.61-1.0) |
|  | 1:160 | 0.92 (0.73-1.0) | 0.0163* |  |  |  |
| All<br>viruses<br>(n=36) | 1:20 | 1.0 (1.0 –1.0) | <0.0001* | 55.04 | 0.83 (0.61-0.94) | 1.0 (0.82-1.0) |
|  | 1:40 | 1.0 (1.0 –1.0) | <0.0001* | 36.36 | 0.83 (0.61-0.94) | 1.0 (0.82-1.0) |
|  | 1:80 | 0.97 (0.92-1.0) | <0.0001* | 17.53 | 0.83 (0.61-0.94) | 0.94 (0.74-0.99) |
|  | 1:160 | 0.84 (0.70-0.98) | 0.0004* |  |  |  |

Median virus neutralization values using liver homogenates in adapted plaque reduction neutralization assays were used for ROC analyses. Samples were grouped (n=12; 6 infected + 6 mock) and compared by dilution. Analyses were made for each virus separately and also considering all viruses combined (all) (n=36). YFV: Yellow fever virus, MAYV: Mayaro virus, CHIKV: Chikungunya virus. AUC: area under the curve. Se: sensitivity, Sp: specificity, dil: dilution. Analyses were carried out with GraphPad Prism 8.0.2. The asterisks indicate statistical significance.

Using a more conservative approach, we ran ROC curves comparing the lowest median neutralization values from the infected groups and the highest median neutralization values from the mock groups (from three independent assays, Table 1). The analyses showed that the YFV-, MAYV- and CHIKV-infected groups were clearly distinguished from the mock groups at dilutions 1:20 and 1:40 (AUC ≥0.92, 95%CI 0.74 – 1.0, p≤0.0163) (S1 Table). When all infected groups were combined and compared to all mock groups, evident differentiation between the groups was observed at dilutions 1:20 and 1:40 (AUC ≥ 0.92; 95% CI ≥ 0.83 – 1.0, p≤ 0.0001) (S1 Table). This aligns with the GLMM results, as the discrepancy between the neutralization values of infected and mock groups is more pronounced at lower dilutions and diminishes with increasing factors, suggesting greater discriminatory power between infected and uninfected groups at lower dilutions.

The PRNT titer, ran with sera, should be calculated based on a 50% or greater reduction in plaque counts to be considered positive. Regrettably, we did not have adequate volumes of sera from experimentally infected mice to run neutralization assays concurrently and assess the level of neutralizing antibodies in sera in comparison to liver homogenates. The decision to refrain from blood harvesting aimed to replicate the natural condition of carcasses, preserving the blood quantity in the liver of experimental animals, and maximizing the chances of detecting neutralizing antibodies in liver homogenates. Given the presence of nonspecific neutralization in all samples, cutoffs were initially estimated to achieve 83% sensitivity and 100% specificity, enhancing the likelihood of identifying true positives and preventing false positives. Targeting 83% of sensitivity and 100% of specificity, for dilution 1:20, cutoffs of 57.96, 56.59, 46.04, and 55.04% neutralization were estimated for YFV, MAYV, CHIKV, and all viruses combined, respectively (Table 5). For dilution 1:40, cutoffs of 40.85, 39.91, 30.13, and 36.36% were estimated for YFV, MAYV, CHIKV, and all viruses combined, respectively (Table 5). We recommend considering a more conservative approach, incorporating a cutoff of 60% neutralization at dilution 1:20 and 40% at dilution 1:40 (sensibility of 0.67 (95%CI 0.44 – 0.84) and specificity of 1.0 (95%CI 0.82 – 1.0), while also observing the proportional reduction in neutralization as the dilution factor increases.

### Investigation of YFV neutralization using liver homogenates from carcasses of free-living neotropical non-human primates

Having established the feasibility of detecting specific virus neutralization activity in liver homogenates from experimentally infected animals, we opted to test liver homogenates obtained from carcasses of free-living NHP collected in rural areas, during the YF outbreak in Minas Gerais, Brazil, in 2017. Frozen liver samples were used to prepare the liver homogenates. Liver homogenates were clarified as described, and an additional step of filtration was added to enhance the clarification process and prevent bacterial contamination from the carcasses. We used 25 samples from NHP carcasses that had undergone prior screening for YFV by RT-qPCR (Sacchetto et al, 2020a), including 8 samples positive for YFV infection confirmed by RT-qPCR (with Cq values in RT-PCR ranging from 8.0 to 35.3) and 17 negative ones (Table 6). The carcasses belonged to animals of different genera, including *Alouatta* sp., *Callicebus* sp., and *Callithrix* sp. that were considered in a good (n=13) or intermediate (n=10), or bad (n=1) preservation status, based on previously established criteria (Sacchetto et al., 2020a).

**Table 6:**
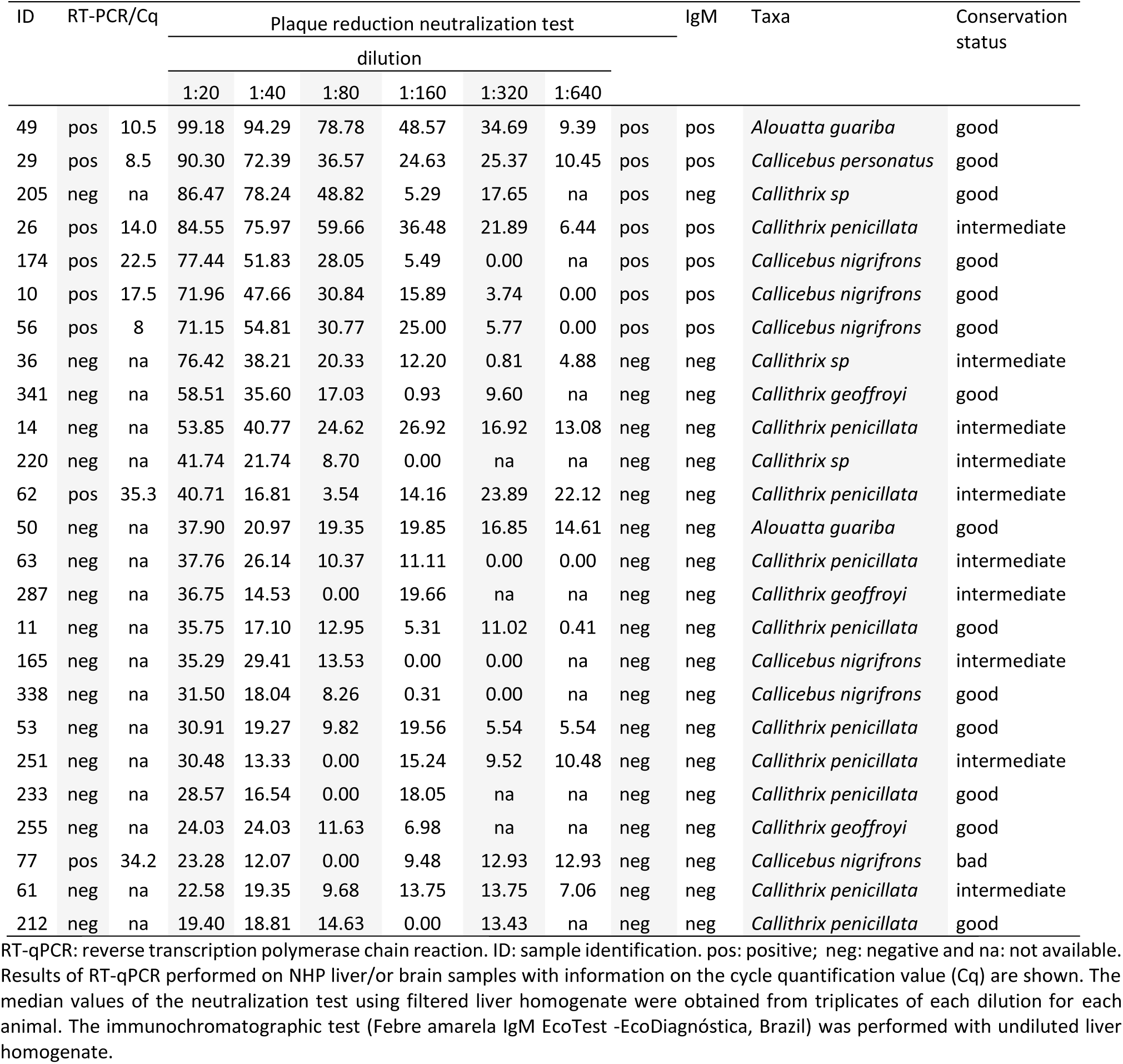
Investigation of antibodies against yellow fever virus using adapted neutralization test, and immunochromatographic tests (IgM) using liver homogenates from non-human primate carcasses.

Twenty-five samples, previously examined using RT-qPCR for the YFV genome, underwent further testing using PRNT, with analysis conducted based on predefined cutoffs and the observing gradual reduction in neutralization proportionally to the increase in dilution factor. Seven samples were positive by the adapted PRNT, as they showed neutralization values ranging from 71.15% to 99.30% at dilution 1:20 and from 47.66% to 94.29% at dilution 1:40. The positive samples in PRNT belonged to specimens of *Callithrix sp, Callicebus sp and Alouatta sp*. (Table 6).

Following the PRNT, undiluted liver homogenates from the 25 samples were subjected to immunochromatographic tests for anti-IgM. All samples yielded positive results in the control test (approximately after 2 minutes the test started). Six out of seven positive samples in PRNT and PCR were positive for IgM (Table 6). Eighteen samples tested negative for the presence of IgM anti-YFV, and were also considered negatives in the PRNT, since they did not present both 60% and 40% cutoffs of neutralization values at dilution 1:20 and 1:40, respectively (Table 6).

These results confirmed the detection of anti-YFV antibodies in the liver homogenate from samples with confirmed infection by YFV. The immunochromatography test used here presents high sensitivity (87.6%) and specificity (98%) according to the manufacturer, which attests to a good probability of differentiating true positive samples from negative ones. Thus, the results obtained from the immunochromatographic tests reinforce the results observed for our PRNT assays, which indicate the presence of antibodies against YFV in liver homogenates. One of the samples (ID 205 – Table 6) presenting high neutralizing activity (from 86.47% to 48.82 at dilutions 1:20 up to 1:80) was negative for YFV by PCR and for IgM. The detection of high neutralization activity without presence of IgM and viral RNA could be linked to later times post-infection, and presence of IgG replacing IgM. Unfortunately, we were not able to test the samples for IgG, using the liver homogenate.

Our findings demonstrate the viability of using liver homogenate from carcasses to investigate antibodies against three different arboviruses using adapted PRNT. Consistent with our results, prior studies have shown the detection of antibodies in solid tissues from experimentally infected animals or blood from human cadavers (Gamble and Patrascu, 1996; Lai et al., 2022). Nevertheless, careful attention must be given when preparing liver homogenate to minimize the presence of particulate organic matter and nonspecific neutralization during PRNT. Samples should undergo clarification by centrifugation until no visible pellet remains, followed by inactivation, re-clarification, and filtration. The results of the adapted neutralization test using liver homogenate should be interpreted considering the appropriate cutoffs per dilution along with the reduction in neutralization according to the dilution. It is crucial to note that even in cases of individuals previously exposed to a particular virus, antibody levels may be low, potentially resulting in false negatives.

Here, we detected antibodies against YFV in liver homogenates of NHPs using rapid tests and PRNT. We observed neutralization activity in samples that were positive or negative by RT-qPCR. During the investigation of outbreaks and epizootics, such as those caused by YFV, molecular or serological tests using serum should be conducted whenever possible to assess infection. However, in some instances serum samples are unavailable or biological samples are unsuitable for molecular testing, leading to inconclusive results. Carcasses are convenient samples that can be used for outbreak investigations, and the potential use of liver homogenates in serological tests expands the possibilities for the investigation of outbreaks and epizootics. A positive result in serological tests, such as the immunochromatographic test or the proposed adapted PRNT can trigger further investigations in specific areas, prompting the collection of new samples to investigate viral infections in animals or humans. Therefore, we propose this approach as a complementary strategy for investigating epizootics, particularly those caused by YFV.

## Acknowledgments

We do thank the Centers for Research in Emerging Infectious Diseases (CREID), CREID Pilot Research Program, the Coordinating Research on Emerging Arboviral Threats Encompassing the Neotropics (CREATE-NEO). We thank Dr. Natália Ingrid Oliveira Silva (currently affiliated to University of Texas Medical Branch) for performing a pilot experiment which launched this project; Dr. Benoit de Thoisy (Institute Pasteur – French Guiana) for initial discussions previously to this work; the Laboratório de Zoonoses, Centro de Controle de Zoonoses da Prefeitura de Belo Horizonte, and Secretaria de Estado de Saúde de Minas Gerais; and lastly colleagues from the Laboratório de Vírus/UFMG, Pró-Reitorias de Graduação, de Pós-graduação, de Pesquisa/Universidade Federal de Minas Gerais/Brazil. We do thank Dr. Mauricio Lacerda Nogueira (FAMERP-Brazil), and Dr. Pedro Augusto Alves for providing virus samples.

## REFERENCES

1. Abreu FVS, Ribeiro IP, Ferreira-de-Brito A, Santos AACD, Miranda RM, Bonelly IS, Neves MSAS, Bersot MI, Santos TPD, Gomes MQ, Silva JLD, Romano APM, Carvalho RG, Said RFDC, Ribeiro MS, Laperrière RDC, Fonseca EOL, Falqueto A, Paupy C, Failloux AB, Moutailler S, Castro MG, Gómez MM, Motta MA, Bonaldo MC, Lourenço-de-Oliveira R. Haemagogus leucocelaenus and Haemagogus janthinomys are the primary vectors in the major yellow fever outbreak in Brazil, 2016-2018. Emerg Microbes Infect. 2019;8(1):218–231. doi: 10.1080/22221751.2019.1568180.

2. Albarnaz JD, De Oliveira LC, Torres AA, Palhares RM, Casteluber MC, Rodrigues CM, Cardozo PL, De Souza AM, Pacca CC, Ferreira PC, Kroon EG, Nogueira ML, Bonjardim CA. MEK/ERK activation plays a decisive role in yellow fever virus replication: implication as an antiviral therapeutic target. Antiviral Res. 2014 Nov;111:82–92. doi: 10.1016/j.antiviral.2014.09.004.

3. Bates, D., Mächler, M., Bolker, B., & Walker, S. (2015). Fitting Linear Mixed-Effects Models Using lme4. Journal of Statistical Software, 67(1), 1–48. 10.18637/jss.v067.i01

4. Brown, Lawrence D., et al. “Interval Estimation for a Binomial Proportion.” Statistical Science, vol. 16, no. 2, 2001, pp. 101–17. JSTOR, http://www.jstor.org/stable/2676784. Accessed 9 Feb. 2024.DIA, Moussa et al. Performance assessment and validation of a plaque reduction neutralization test (PRNT) in support to yellow fever diagnostic and vaccine clinical trials. Journal of Medical Virology, [S. l.], v. 95, n. 4, 2023. ISSN: 0146-6615. DOI: 10.1002/jmv.28700. Disponível em: https://onlinelibrary.wiley.com/doi/10.1002/jmv.28700.

5. Caicedo EY, Charniga K, Rueda A, Dorigatti I, Mendez Y, Hamlet A, Carrera JP, Cucunubá ZM. The epidemiology of Mayaro virus in the Americas: A systematic review and key parameter estimates for outbreak modelling. PLoS Negl Trop Dis. 2021 Jun 3;15(6):e0009418. doi: 10.1371/journal.pntd.0009418. Erratum in: PLoS Negl Trop Dis. 2023 Jan 4;17(1):e0011034.

6. Crawley, Michael J. The R Book. Chichester, West Sussex, United Kingdom: Wiley, 2013.

7. De Azevedo Fernandes NCC, Guerra JM, Díaz-Delgado J, Cunha MS, Saad LD, Iglezias SD, Ressio RA, Dos Santos Cirqueira C, Kanamura CT, Jesus IP, Maeda AY, Vasami FGS, de Carvalho J, de Araújo LJT, de Souza RP, Nogueira JS, Spinola RMF, Catão-Dias JL. Differential Yellow Fever Susceptibility in New World Nonhuman Primates, Comparison with Humans, and Implications for Surveillance. Emerg Infect Dis. 2021 Jan;27(1):47–56. doi: 10.3201/eid2701.191220.

8. Delatorre E, de Abreu FVS, Ribeiro IP, Gómez MM, Dos Santos AAC, Ferreira-de-Brito A, Neves MSAS, Bonelly I, de Miranda RM, Furtado ND, Raphael LMS, da Silva LFF, de Castro MG, Ramos DG, Romano APM, Kallás EG, Vicente ACP, Bello G, Lourenço-de-Oliveira R, Bonaldo MC. Distinct YFV Lineages Co-circulated in the Central-Western and Southeastern Brazilian Regions From 2015 to 2018. Front Microbiol. 2019 May 24;10:1079. doi: 10.3389/fmicb.2019.01079.

9. Dia M, Bob NS, Talla C, Dupressoir A, Escadafal C, Thiam MS, Diallo A, Ndiaye O, Heraud JM, Faye O, Sall AA, Faye O, Fall G. Performance assessment and validation of a plaque reduction neutralization test (PRNT) in support to yellow fever diagnostic and vaccine clinical trials. J Med Virol. 2023 Apr;95(4):e28700. doi: 10.1002/jmv.28700.

10. Diagne CT, Bengue M, Choumet V, Hamel R, Pompon J, Missé D. Mayaro Virus Pathogenesis and Transmission Mechanisms. Pathogens. 2020 Sep 8;9(9):738. doi: 10.3390/pathogens9090738.

11. Domingo C, Patel P, Yillah J, Weidmann M, Méndez JA, Nakouné ER, Niedrig M. Advanced yellow fever virus genome detection in point-of-care facilities and reference laboratories. J Clin Microbiol. 2012 Dec;50(12):4054–60. doi: 10.1128/JCM.01799-12. Epub 2012 Oct 10. PMID: 23052311; PMCID: PMC3503008.

12. Edwards CJ, Welch SR, Chamberlain J, Hewson R, Tolley H, Cane PA, Lloyd G. Molecular diagnosis and analysis of Chikungunya virus. J Clin Virol. 2007 Aug;39(4):271–5. doi: 10.1016/j.jcv.2007.05.008.

13. Erickson AK, Pfeiffer JK. Dynamic viral dissemination in mice infected with yellow fever virus strain 17D. J Virol. 2013 Nov;87(22):12392–7. doi: 10.1128/JVI.02149-13.

14. Figueiredo PO, Stoffella-Dutra AG, Barbosa Costa G, Silva de Oliveira J, Dourado Amaral C, Duarte Santos J, Soares Rocha KL, Araújo Júnior JP, Lacerda Nogueira M, Zazá Borges MA, Pereira Paglia A, Desiree LaBeaud A, Santos Abrahão J, Geessien Kroon E, Bretas de Oliveira D, Paiva Drumond B, de Souza Trindade G. Re-Emergence of Yellow Fever in Brazil during 2016-2019: Challenges, Lessons Learned, and Perspectives. Viruses. 2020 Oct 30;12(11):1233. doi: 10.3390/v12111233.

15. Fumagalli MJ, de Souza WM, de Castro-Jorge LA, de Carvalho RVH, Castro ÍA, de Almeida LGN, Consonni SR, Zamboni DS, Figueiredo LTM. Chikungunya Virus Exposure Partially Cross-Protects against Mayaro Virus Infection in Mice. J Virol. 2021 Nov 9;95(23):e0112221. doi: 10.1128/JVI.01122-21.

16. Gamble HR, Patrascu IV. Whole Blood, Serum, and Tissue Fluids in an Enzyme Immunoassay for Swine Trichinellosis. J Food Prot. 1996 Nov;59(11):1213–1217. doi: 10.4315/0362-028X-59.11.1213.

17. Gibney KB, Edupuganti S, Panella AJ, Kosoy OI, Delorey MJ, Lanciotti RS, Mulligan MJ, Fischer M, Staples JE. Detection of anti-yellow fever virus immunoglobulin m antibodies at 3-4 years following yellow fever vaccination. Am J Trop Med Hyg. 2012 Dec;87(6):1112–5. doi: 10.4269/ajtmh.2012.12-0182.

18. Giovanetti M, de Mendonça MCL, Fonseca V, Mares-Guia MA, Fabri A, Xavier J, de Jesus JG, Gräf T, Dos Santos Rodrigues CD, Dos Santos CC, Sampaio SA, Chalhoub FLL, de Bruycker Nogueira F, Theze J, Romano APM, Ramos DG, de Abreu AL, Oliveira WK, do Carmo Said RF, de Alburque CFC, de Oliveira T, Fernandes CA, Aguiar SF, Chieppe A, Sequeira PC, Faria NR, Cunha RV, Alcantara LCJ, de Filippis AMB. Yellow Fever Virus Reemergence and Spread in Southeast Brazil, 2016-2019. J Virol. 2019 Dec 12;94(1):e01623–19. doi: 10.1128/JVI.01623-19. Erratum in: J Virol. 2020 May 18;94(11):

19. Giovanetti M, Pinotti F, Zanluca C, Fonseca V, Nakase T, Koishi AC, Tscha M, Soares G, Dorl GG, Marques AEML, Sousa R, Adelino TER, Xavier J, de Oliveira C, Patroca S, Guimaraes NR, Fritsch H, Mares-Guia MA, Levy F, Passos PH, da Silva VL, Pereira LA, Mendonça AF, de Macêdo IL, Ribeiro de Sousa DE, Rodrigues de Toledo Costa G, Botelho de Castro M, de Souza Andrade M, de Abreu FVS, Campos FS, Iani FCM, Pereira MA, Cavalcante KRLJ, de Freitas ARR, Campelo de Albuquerque CF, Macário EM, Dos Anjos MPD, Ramos RC, Campos AAS, Pinter A, Chame M, Abdalla L, Riediger IN, Ribeiro SP, Bento AI, de Oliveira T, Freitas C, Oliveira de Moura NF, Fabri A, Dos Santos Rodrigues CD, Dos Santos CC, Barreto de Almeida MA, Dos Santos E, Cardoso J, Augusto DA, Krempser E, Mucci LF, Gatti RR, Cardoso SF, Fuck JAB, Lopes MGD, Belmonte IL, Mayoral Pedroso da Silva G, Soares MRF, de Castilhos MMS, de Souza E Silva JC, Bisetto Junior A, Pouzato EG, Tanabe LS, Arita DA, Matsuo R, Dos Santos Raymundo J, Silva PCL, Santana Araújo Ferreira Silva A, Samila S, Carvalho G, Stabeli R, Navegantes W, Moreira LA, Ferreira AGA, Pinheiro GG, Nunes BTD, de Almeida Medeiros DB, Cruz ACR, Venâncio da Cunha R, Van Voorhis W, Bispo de Filippis AM, Almiron M, Holmes EC, Ramos DG, Romano A, Lourenço J, Alcantara LCJ, Duarte Dos Santos CN. Genomic epidemiology unveils the dynamics and spatial corridor behind the Yellow Fever virus outbreak in Southern Brazil. Sci Adv. 2023 Sep;9(35):eadg9204. doi: 10.1126/sciadv.adg9204.

20. Hallengärd D, Kakoulidou M, Lulla A, Kümmerer BM, Johansson DX, Mutso M, Lulla V, Fazakerley JK, Roques P, Le Grand R, Merits A, Liljeström P. Novel attenuated Chikungunya vaccine candidates elicit protective immunity in C57BL/6 mice. J Virol. 2014 Mar;88(5):2858–66. doi: 10.1128/JVI.03453-13.

21. Hothorn T, Bretz F, Westfall P. Simultaneous Inference in General Parametric Models. Biometrical Journal. 2008. 50(3), 346–363.

22. Kading RC, Borland EM, Cranfield M, Powers AM. Prevalence of antibodies to alphaviruses and flaviviruses in free-ranging game animals and nonhuman primates in the greater Congo basin. J Wildl Dis. 2013 Jul;49(3):587–99. doi: 10.7589/2012-08-212.

23. Lai A, Tambuzzi S, Bergna A, Battistini A, Della Ventura C, Galli M, Zoja R, Zehender G, Cattaneo C. Evidence of SARS-CoV-2 Antibodies and RNA on Autopsy Cases in the Pre-Pandemic Period in Milan (Italy). Front Microbiol. 2022 Jun 15;13:886317. doi: 10.3389/fmicb.2022.886317.

24. Lautt, W. Wayne. Hepatic Circulation. Colloquium Series on Integrated Systems Physiology: From Molecule to Function, [S. l.], v. 1, n. 1, p. 1–174, 2009. ISSN: 2154560X. DOI: 10.4199/C00004ED1V01Y200910ISP001. Disponível em: https://access.portico.org/stable?au=pgg3gvb7b5g.

25. Lindsey NP, Horiuchi KA, Fulton C, Panella AJ, Kosoy OI, Velez JO, Krow-Lucal ER, Fischer M, Staples JE. Persistence of yellow fever virus-specific neutralizing antibodies after vaccination among US travellers. J Travel Med. 2018 Jan 1;25(1):10.1093/jtm/tay108. doi: 10.1093/jtm/tay108.

26. Ministério da Saúde/Brasil. Secretaria de Vigilância em Saúde. Departamento de Imunização e Doenças Transmissíveis. 2021 PLANO DE CONTINGÊNCIA PARA RESPOSTA ÀS EMERGÊNCIAS EM SAÚDE PÚBLICA FEBRE AMARELA. 2ª edição. Disponível em https://www.gov.br/saude/pt-br/assuntos/saude-de-a-a-z/f/febre-amarela/publicacoes/plano_contingencia_emergencias_febre_amarela_2_ed.pdf. (acessed in August 09 2023).

27. Ministério da Saúde/Brasil. Secretaria de Vigilância em Saúde. Departamento de Imunização e Doenças Transmissíveis. 2017. GUIA DE VIGILÂNCIA DE EPIZOOTIAS EM PRIMATAS NÃO HUMANOS E ENTOMOLOGIA APLICADA À VIGILÂNCIA DA FEBRE AMARELA. http://www.vs.saude.ms.gov.br/wp-content/uploads/2018/01/Guia_Epizootias_Febre_Amarela_2a_ed_atualizada_2017.pdf (acessed in August 09 2023).

28. Ministério da Saúde/Brasil. Secretaria de Vigilância em Saúde. Departamento de Imunização e Doenças Transmissíveis. 2020. Manual de Manejo Clínico da Febre amarela. Disponível em https://bvsms.saude.gov.br/bvs/publicacoes/manual_manejo_clinico_febre_amarela.pdf. (acessed in August 09 2023).

29. Ministério da Saúde/Brasil, Secretaria de Atenção à Saúde. 2017. Febre amarela: guia para profissionais de saúde. Disponível em https://bvsms.saude.gov.br/bvs/publicacoes/febre_amarela_guia_profissionais_saude.pdf. (acessed in August 07 2023).

30. Ministério da Saúde/Brasil, Secretaria de Vigilância em Saúde. 2015. Reemergência da Febre Amarela Silvestre no Brasil, 2014/2015: situação epidemiológica e a importância da vacinação preventiva e da vigilância intensificada no período sazonal. Boletim Epidemiológico.

31. Moreira-Soto A, Carneiro IO, Fischer C, Feldmann M, Kümmerer BM, Silva NS, Santos UG, Souza BFCD, Liborio FA, Valença-Montenegro MM, Laroque PO, da Fontoura FR, Oliveira AVD, Drosten C, de Lamballerie X, Franke CR, Drexler JF. Limited Evidence for Infection of Urban and Peri-urban Nonhuman Primates with Zika and Chikungunya Viruses in Brazil. mSphere. 2018 Jan 31;3(1):e00523–17. doi: 10.1128/mSphere.00523-17.

32. Moreno ES, Spinola R, Tengan CH, Brasil RA, Siciliano MM, Coimbra TL, Silveira VR, Rocco IM, Bisordi I, Souza RP, Petrella S, Pereira LE, Maeda AY, Silva FG, Suzuki A. Yellow fever epizootics in non-human primates, São Paulo state, Brazil, 2008-2009. Rev Inst Med Trop Sao Paulo. 2013 Jan-Feb;55(1):45–50. doi: 10.1590/s0036-46652013000100008.

33. Mota MTO, Costa VV, Sugimoto MA, Guimarães GF, Queiroz-Junior CM, Moreira TP, de Sousa CD, Santos FM, Queiroz VF, Passos I, Hubner J, Souza DG, Weaver SC, Teixeira MM, Nogueira ML. In-depth characterization of a novel live-attenuated Mayaro virus vaccine candidate using an immunocompetent mouse model of Mayaro disease. Sci Rep. 2020 Mar 24;10(1):5306. doi: 10.1038/s41598-020-62084-x.

34. Narat V, Kampo M, Heyer T, Rupp S, Ambata P, Njouom R, Giles-Vernick T. Using physical contact heterogeneity and frequency to characterize dynamics of human exposure to nonhuman primate bodily fluids in central Africa. PLoS Negl Trop Dis. 2018 Dec 27;12(12):e0006976. doi: 10.1371/journal.pntd.0006976.

35. Naveca FG, Nascimento VAD, Souza VC, Nunes BTD, Rodrigues DSG, Vasconcelos PFDC. Multiplexed reverse transcription real-time polymerase chain reaction for simultaneous detection of Mayaro, Oropouche, and Oropouche-like viruses. Mem Inst Oswaldo Cruz. 2017 Jul;112(7):510–513. doi: 10.1590/0074-02760160062.

36. Pan American Health Organization (PAHO). Febre amarela. 2023. Disponível em https://www.paho.org/pt/topicos/febre-amarela. (acesso 06 de agosto de 2023).

37. Pan American Health Organization (PAHO). Alerta Epidemiológico: Febre Amarela. 31 de agosto de 2022, Washington, D.C.: OPAS/OMS; 2022. Disponível em https://www.paho.org/pt/documentos/alerta-epidemiologico-febre-amarela-31-agosto-2022 (acessed in August 09 2023).

38. Possas C, Lourenço-de-Oliveira R, Tauil PL, Pinheiro FP, Pissinatti A, Cunha RVD, Freire M, Martins RM, Homma A. Yellow fever outbreak in Brazil: the puzzle of rapid viral spread and challenges for immunisation. Mem Inst Oswaldo Cruz. 2018 Sep 3;113(10):e180278. doi: 10.1590/0074-02760180278.

39. Reinhardt, B., Jaspert, R., Niedrig, M., Kostner, C. and L’age-Stehr, J. (1998), Development of viremia and humoral and cellular parameters of immune activation after vaccination with yellow fever virus strain 17D: A model of human flavivirus infection. J. Med. Virol., 56: 159–167. 10.1002/(SICI)1096-9071(199810)56:2<159::AID-JMV10>3.0.CO;2-B

40. Rezende IM, Sacchetto L, Munhoz de Mello É, Alves PA, Iani FCM, Adelino TÉR, Duarte MM, Cury ALF, Bernardes AFL, Santos TA, Pereira LS, Dutra MRT, Ramalho DB, de Thoisy B, Kroon EG, Trindade GS, Drumond BP. Persistence of Yellow fever virus outside the Amazon Basin, causing epidemics in Southeast Brazil, from 2016 to 2018. PLoS Negl Trop Dis. 2018 Jun 4;12(6):e0006538. doi: 10.1371/journal.pntd.0006538.

41. R Core Team (2023) - R Core Team (2023). R: A Language and Environment for Statistical Computing. R Foundation for Statistical Computing, Vienna, Austria. <https://www.R-project.org/>.

42. Sacchetto L, Silva NIO, Rezende IM, Arruda MS, Costa TA, de Mello ÉM, Oliveira GFG, Alves PA, de Mendonça VE, Stumpp RGAV, Prado AIA, Paglia AP, Perini FA, Lacerda Nogueira M, Kroon EG, de Thoisy B, Trindade GS, Drumond BP. Neighbor danger: Yellow fever virus epizootics in urban and urban-rural transition areas of Minas Gerais state, during 2017-2018 yellow fever outbreaks in Brazil. PLoS Negl Trop Dis. 2020a Oct 5;14(10):e0008658. doi: 10.1371/journal.pntd.0008658.

43. Sacchetto L, Drumond BP, Han BA, Nogueira ML, Vasilakis N. Re-emergence of yellow fever in the neotropics - quo vadis? Emerg Top Life Sci. 2020b Dec 11;4(4):399–410. doi: 10.1042/ETLS20200187.

44. Silva NIO, Albery GF, Arruda MS, Oliveira GFG, Costa TA, de Mello ÉM, Moreira GD, Reis EV, Silva SAD, Silva MC, de Almeida MG, Becker DJ, Carlson CJ, Vasilakis N, Hanley KA, Drumond BP. Ecological drivers of sustained enzootic yellow fever virus transmission in Brazil, 2017-2021. PLoS Negl Trop Dis. 2023 Jun 5;17(6):e0011407. doi: 10.1371/journal.pntd.0011407.

45. Silva NIO, Sacchetto L, de Rezende IM, Trindade GS, LaBeaud AD, de Thoisy B, Drumond BP. Recent sylvatic yellow fever virus transmission in Brazil: the news from an old disease. Virol J. 2020 Jan 23;17(1):9. doi: 10.1186/s12985-019-1277-7.

46. Terzian, A.C.B., Zini, N., Sacchetto, L. et al. Evidence of natural Zika virus infection in neotropical non-human primates in Brazil. Sci Rep 8, 16034 (2018). 10.1038/s41598-018-34423-6

47. Timiryasova TM, Bonaparte MI, Luo P, Zedar R, Hu BT, Hildreth SW. Optimization and validation of a plaque reduction neutralization test for the detection of neutralizing antibodies to four serotypes of dengue virus used in support of dengue vaccine development. Am J Trop Med Hyg. 2013 May;88(5):962–970. doi: 10.4269/ajtmh.12-0461.

48. Valentine MJ, Murdock CC, Kelly PJ. Sylvatic cycles of arboviruses in non-human primates. Parasit Vectors. 2019 Oct 2;12(1):463. doi: 10.1186/s13071-019-3732-0.

49. Waggoner JJ, Rojas A, Pinsky BA. Yellow Fever Virus: Diagnostics for a Persistent Arboviral Threat. J Clin Microbiol. 2018 Sep 25;56(10):e00827–18. doi: 10.1128/JCM.00827-18.

50. Wickham, H. ggplot2: Elegant Graphics for Data Analysis. [s.l.] : Springer-Verlag New York, 2016. ISBN: 978-3-319-24275-0.

51. Wilson, Edwin B. “Probable Inference, the Law of Succession, and Statistical Inference.” Journal of the American Statistical Association, vol. 22, no. 158, 1927, pp. 209–12. JSTOR, 10.2307/2276774. Accessed 9 Feb. 2024.

52. World Health Organization (WHO). Yellow fever. 2024. Available at https://www.who.int/teams/health-product-policy-and-standards/standards-and-specifications/vaccines-quality/yellow-fever. Accessed in 03-27-2024.

