## Supplementary material for "Detection of neutralizing antibodies against arboviruses from liver homogenates": Suplementary material

Costa et al.

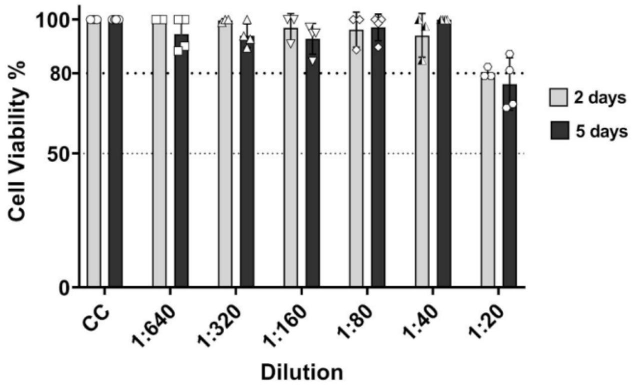


**S1 Figure. Cytotoxicity by MTT assay in C57BL/6 and IFNAR$^{-}$/$^{-}$ animals.** Liver homogenates of C57BL/6 and IFNAR$^{-}$/$^{-}$ animals were incubated in Vero cells CCL-81 for two and five days, respectively. Then, the MTT cytotoxicity was evaluated by absorbance reading. For two days of incubation (grey bars), the mean cell viability was equivalent to 80.13% at dilution 1:20 and over 80% in higher dilutions. For five days of incubation (black bars), the mean cell viability was 75.97% at dilution 1:20 and over 80% at higher dilutions.

A

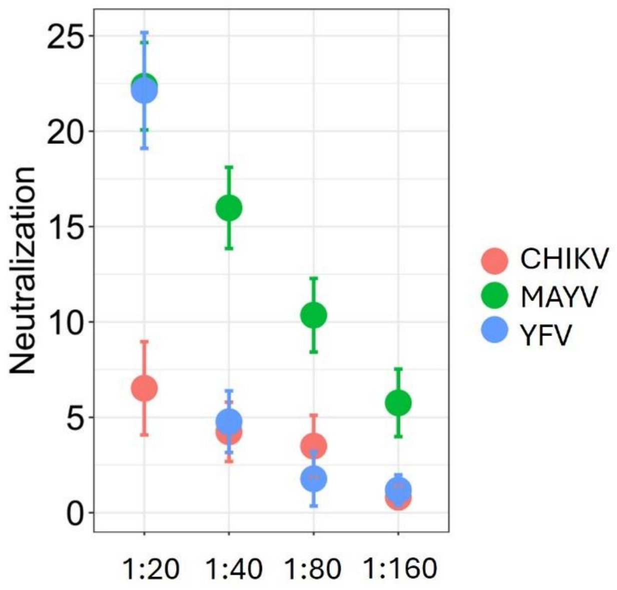


B

| virus group/dilution | | Mock MAYV 1:20 | Mock MAYV 1:40 | Mock MAYV 1:80 | Mock MAYV 1:160 | Mock CHIKV 1:20 | Mock CHIKV 1:40 | Mock CHIKV 1:80 | Mock CHIKV 1:160 | Mock YFV 1:40 | Mock YFV 1:80 | Mock YFV 1:160 |
|---|---|---|---|---|---|---|---|---|---|---|---|---|
| Mock YFV | 1:20 | **1.0** | 0.66 | <0.01* | <0.01* | **<0.01*** | <0.01* | <0.01* | <0.01* | <0.01* | <0.01* | <0.01* |
| | 1:40 | <0.01* | **<0.01*** | 0.73 | 1.0 | 1.0 | **1.0** | 1.0 | 0.97 | | 0.99 | 0.99 |
| | 1:80 | <0.01* | <0.01* | **0.03*** | 0.97 | 0.90 | 1.0 | **1.0** | 1.0 | | | 1.0 |
| | 1:160 | <0.01* | <0.01* | 0.07 | **0.79** | 0.80 | 1.0 | 1.0 | **1.0** | | | |
| Mock CHIKV | 1:20 | **<0.01*** | 0.05 | 0.98 | 1.0 | | 1.0 | 1.0 | 0.71 | | | |
| | 1:40 | <0.01* | **<0.01*** | 0.62 | 1.0 | | | 1.0 | 0.99 | | | |
| | 1:80 | <0.01* | <0.01* | **0.20** | 1.0 | | | | 1.0 | | | |
| | 1:160 | <0.01* | <0.01* | 0.05 | **0.70** | | | | | | | |
| Mock MAYV | 1:20 | | 0.56 | <0.01* | <0.01* | | | | | | | |
| | 1:40 | | | 0.74 | 0.02* | | | | | | | |
| | 1:80 | | | | 0.92 | | | | | | | |

**S2 Figure.** (A) Estimates of average median viral neutralization observed in mock groups tested against YFV, MAYV and CHIKV at dilutions 1:20 to 1:160. YFV-, MAYV-, and CHIKV- mock groups consisted of six mice each. (B) Results of post-hoc analyses of generalized linear mixed models, with Tukey correction. The asterisks indicate statistical significance, and the bold numbers indicate the comparison between groups in the respective dilution. YFV: Yellow fever virus (in blue), MAYV: Mayaro virus (in green), CHIKV: Chikungunya virus (in orange). The asterisks indicate statistical significance.

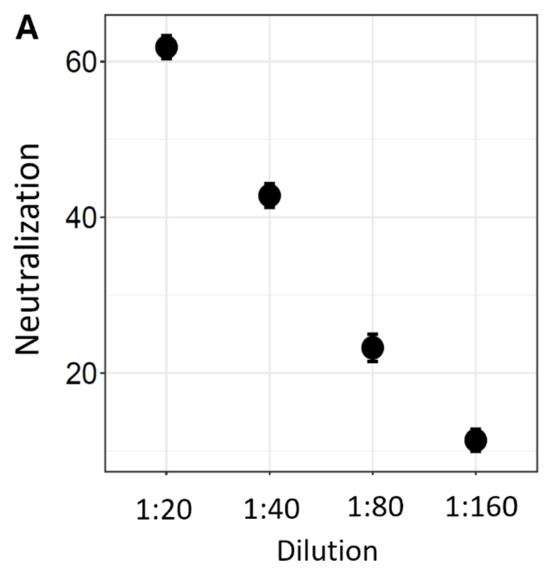


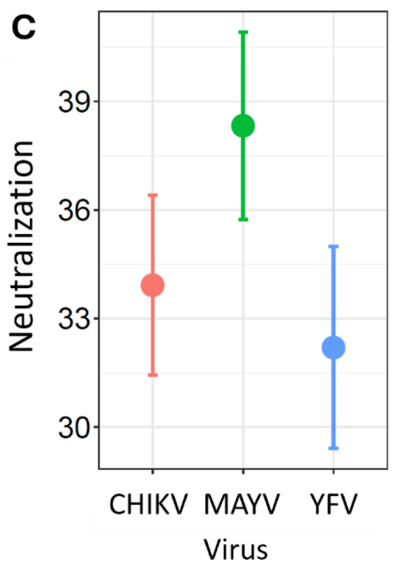


**B**

| Dilution | 1:20 | 1:40 | 1:80 |
|---|---|---|---|
| **1:40** | $<1.0^{-6}$* | | |
| **1:80** | $<1.0^{-6}$* | $<1.0^{-6}$* | |
| **1:160** | $<1.0^{-6}$* | $<1.0^{-6}$* | $<1.0^{-6}$* |

**D**

| Virus/infected group | MAYV | CHIKV |
|---|---|---|
| **YFV** | <0.01* | 0.62 |
| **CHIKV** | 0.044* | |

**S3 Figure.** Comparison of average median virus neutralization in adapted plaque reduction neutralization tests using liver homogenates of YFV-, MAYV-, and CHIKV-infected mice. (A) and (C) Estimates of average median viral neutralization observed in relation to dilution of virus, respectively. (B) and (D) Results of post-hoc analyses of generalized linear mixed models, with Tukey correction. YFV-, MAYV-, and CHIKV- infected groups consisted of six mice each. YFV: Yellow fever virus (in blue), MAYV: Mayaro virus (in green), CHIKV: Chikungunya virus (in orange). The asterisks indicate statistical significance.

**S1 Table:** Receiver operating characteristic (ROC) curve analyses for the lowest median neutralization values of YFV-, MAYV, CHIKV-infected groups compared to the highest median neutralization values of respective mock groups.

| | | Lowest median neutralization values of infected group X Highest median neutralization values of mock group | |
|---|---|---|---|
| Virus (sample size) | dil | AUC (95% CI) | p value |
| YFV (12) | 1:20 | 1.0 (1.0-1.0) | 0.0039* |
| | 1:40 | 1.0 (1.0-1.0) | 0.0039* |
| | 1:80 | 0.64 (0.31-0.97) | 0.4233 |
| | 1:160 | 0.75 (0.46-1.0) | 0.1495 |
| MAYV (12) | 1:20 | 1.0 (1.0-1.0) | 0.0039* |
| | 1:40 | 0.94 (0.81-1) | 0.0104* |
| | 1:80 | 0.58 (0.24 – 0.93) | 0.6310 |
| | 1:160 | 0.75 (0.44-1) | 0.1495 |
| CHIKV (12) | 1:20 | 1.0 (1.0-1.0) | 0.0039* |
| | 1:40 | 0.92 (0.74-1) | 0.0163* |
| | 1:80 | 0.69 (0.38-1) | 0.2623 |
| | 1:160 | 0.64 (0.31 – 0.97) | 0.4233 |
| ALL (36) | 1:20 | 1.0 (1.0-1.0) | <0.0001* |
| | 1:40 | 0.92 (0.83-1.0) | <0.0001* |
| | 1:80 | 0.52 (0.33-0.71) | 0.8247 |
| | 1:160 | 0.59 (0.40-0.78) | 0.3425 |

Lowest (for infected groups) and highest (for mock groups) median neutralization values using liver homogenates in adapted plaque reduction neutralization assays were used for ROC analyses. Samples were grouped (n=12; 6 infected + 6 mock) and compared by dilution. Analyses were made for each virus separately and also considering all data together (n=36). YFV: Yellow fever virus, MAYV: Mayaro virus, CHIKV: Chikungunya virus. AUC: area under the curve. dil: dilution. Analyses were carried out with Graph PadPrism 8.0.2. The asterisks indicate statistical significance.
